# SmbHLH37 functions antagonistically with SmMYC2 in regulating jasmonate-mediated biosynthesis of phenolic acids in *Salvia miltiorrhiza*

**DOI:** 10.1101/438143

**Authors:** Tang-Zhi Du, Jun-Feng Niu, Jiao Su, Sha-Sha Li, Xiao-Rong Guo, Lin Li, Xiao-Yan Cao, Jie-Fang Kang

## Abstract

Jasmonates (JAs) are integral to various defense responses and induce biosynthesis of many secondary metabolites. MYC2, a basic helix-loop-helix (bHLH) transcription factor (TF), acts as a transcriptional activator of JA signaling. MYC2 is repressed by the JASMONATE ZIM-domain (JAZ) proteins in the absence of JA, but de-repressed by the protein complex SCF^COI1^ on perception of JA. We previously reported that overexpression of *SmMYC2* promotes the production of salvianolic acid B (Sal B) in *Salvia miltiorrhiza.* However, the responsible molecular mechanism is unclear. Here, we showed that SmMYC2 binds to and activates the promoters of its target genes *SmTAT1*, *SmPAL1*, and *SmCYP98A14* to activate Sal B accumulations. *SmbHLH37*, a novel bHLH gene significantly up-regulated by constitutive expression of *SmMYC2*, was isolated from *S. miltiorrhiza* for detailed functional characterization. SmbHLH37 forms a homodimer and interacts with SmJAZ3/8. Overexpression of *SmbHLH37* substantially decreased yields of Sal B. SmbHLH37 binds to the promoters of its target genes *SmTAT1* and *SmPAL1* and blocks their expression to suppress the pathway for Sal B biosynthesis. These results indicate that SmbHLH37 negatively regulates JA signaling and functions antagonistically with SmMYC2 in regulating Sal B biosynthesis in *S. miltiorrhiza*.

## Introduction

*Salvia miltiorrhiza* Bunge, a well-known member of the Labiatae family, is considered a model medicinal plant (Guo *et al.*, 2014). Its dry roots and rhizomes (called ‘danshen' in Chinese) are widely applied in the treatment of various cerebrovascular and cardiovascular diseases (Han *et al.*, 2008; Zeng *et al.*, 2013; Su *et al.*, 2015). The major bioactive components of *S. miltiorrhiza* are classified as water-soluble phenolic acids, including salvianolic acid B (Sal B) and rosmarinic acid (RA); and lipid-soluble tanshinones such as cryptotanshinone and tanshinone IIA (Ma *et al.*, 2013; Ma *et al.*, 2015). Phenolic acids are attracting increased attention because of their marked pharmacological activities coupled with their traditional use as herbs steeped in boiling water in China. Among these phenolic acids, Sal B is predominant and is regarded for its antioxidant properties and scavenging of free radicals (Zhao *et al.*, 2008). It offers protection against fibrosis, tumor development, aging, and cardiovascular/cerebrovascular diseases (Zhao *et al.*, 2008; Tsai *et al.*, 2010).

The biosynthetic pathway of Sal B consists of a phenylalanine-derived pathway and tyrosine-derived pathway (Di *et al.*, 2013; Ma *et al.*, 2013; Wang *et al.*, 2015). In view of the economic value and clinical demand for this active ingredient, biological approaches have been taken to augment its synthesis, including the engineering of genes in the biosynthetic pathway and ectopic expression of transcription factors (TFs) (Zhang *et al.*, 2010; Wang *et al.*, 2013; Zhang *et al.*, 2014; Zhou *et al.*, 2016; Yang *et al.*, 2017). For example, AtPAP1 from *Arabidopsis thaliana* is a transcriptional activator of phenolic acid biosynthesis in *S. miltiorrhiza* (Zhang *et al.*, 2010; Zhang *et al.*, 2014). Heterologous expression of two transcription factors, Delila (DEL) and Rosea1 (ROS1) from *Antirrhinum majus*, significantly elevates the production of Sal B in *S. miltiorrhiza* (Wang *et al.*, 2013). In addition, exogenous application of methyl jasmonate (MeJA) triggers an extensive transcriptional reprogramming of metabolism and dramatically increases Sal B biosynthesis in that species (Ge *et al.*, 2015).

Jasmonates (JAs) play crucial roles in plant responses to various stimuli and induce biosynthesis of many secondary metabolites (Browse, 2009; Zhou and Memelink, 2016). The Jasmonate ZIM-domain (JAZ) proteins function as negative regulators to repress diverse JA responses (Chini *et al.*, 2007; Thines *et al.*, 2007; Seo *et al.*, 2011; Song *et al.*, 2011). Jasmonoyl-_L_-isoleucine (JA-Ile), the active form of JA, promotes the degradation of Jasmonate ZIM-domain (JAZ) proteins via the 26S proteasome system (Farmer, 2007; Sheard *et al.*, 2010). This is followed by de-repression of MYC2, a basic helix-loop-helix (bHLH) TF that has a central role in JA signaling, resulting in transcriptional activation of downstream target genes (Lorenzo *et al.*, 2004; Chico *et al.*, 2008; Katsir *et al.*, 2008). Nine JAZ genes have been cloned from *S. miltiorrhiza* and some have been functionally verified as negative regulators of active ingredients in this species. For example, overexpression of *SmJAZ8* de-regulates the yields of salvianolic acids and tanshinones in MeJA-induced transgenic hairy roots (Ge *et al.*, 2015; Pei *et al.*, 2018). Both SmJAZ3 and SmJAZ9 act as repressive transcriptional regulators in the biosynthesis of tanshinones (Shi *et al.*, 2016). However, SmMYC2a and SmMYC2b, two orthologs of MYC2, interact with SmJAZs and positively regulate the biosynthesis of tanshinones and Sal B in *S. miltiorrhiza* hairy roots (Zhou *et al.*, 2016).

The bHLH proteins, one of the largest TF families in plants, modulate various physiological or morphological events, including different branches of the flavonoid pathway (Carretero-Paulet *et al.*, 2010; Hichri *et al.*, 2011). The bHLH family consists of an N-terminal stretch of basic amino acid residues responsible for DNA binding and an HLH domain to form homo-or heterodimers (Goossens *et al.*, 2016), which bind E-box sequences (CANNTG), such as the G-box (CACGTG), in the promoter of their target genes (Ezer *et al.*, 2017). The bHLHs are monophyletic and constitute 26 subfamilies characterized by the presence of highly conserved short amino acid motifs (Pires and Dolan, 2010). MYC2, a member of bHLH subgroup IIIe, positively regulates secondary metabolism during JA signaling in a species-specific manner (Dombrecht *et al.*, 2007; Todd *et al.*, 2010; Zhang *et al.*, 2011). JA-ASSOCIATED MYC2-LIKE1 (JAM1), JAM2, and JAM3 (bHLH17, −13, and −3, respectively) belong to the bHLH IIId subfamily in *A. thaliana*. Each contains a domain that can interact with JAZ proteins and negatively regulate JA responses (Fonseca *et al.*, 2014; Sasaki-Sekimoto *et al.*, 2014). JAM1 substantially reduces those responses, inhibiting root growth and interrupting anthocyanin accumulations and male fertility (Nakata *et al.*, 2013). JAM2 and JAM3 have the same functions and act redundantly with JAM1 (Nakata and Ohme-Takagi, 2013). These JAMs antagonize MYC2, MYC3, and MYC4 during JA-induced leaf senescence by binding to the same target sequences of MYC-activated genes (Qi *et al.*, 2015a).

Zhang *et al.*, (2015) have identified 127 bHLH genes in *S. miltiorrhiza* based on genome-wide analyses. They have predicted seven *bHLH*s, including *SmbHLH37*, that are involved in tanshinone biosynthesis. However, the functions of those genes have not been characterized. We previously reported that overexpression of *SmMYC2* increases the production of phenolic acids in *S. miltiorrhiza* (Yang *et al.*, 2017). Further investigation showed that constitutive expression of that gene significantly up-regulates transcript levels of *SmbHLH37* (Su *et al.*, 2017). Multiple alignments of the SmbHLH37 protein sequence with AtbHLHs from *Arabidopsis* have indicated that SmbHLH37 is most closely correlated with AtbHLH3 (JAM3), both of which belong to the IIId subfamily (Su *et al.*, 2017). In the present study, we identified SmbHLH37 as a new target of JAZ proteins. We then conducted overexpression experiments to explore the function of *SmbHLH37* in *S. miltiorrhiza*. Transgenic overexpressing (OE) plants showed significantly lower accumulations of Sal B. We concluded that SmbHLH37 antagonizes the previously reported transcription activator SmMYC2 in controlling salvianolic acid biosynthesis in *S. miltiorrhiza* by binding to their downstream target sequences. Coordinated regulation of Sal B by this transcription repressor and activator provides clues about the previously unknown complex mechanism for directing the production of secondary metabolites.

## Materials and methods

### Experimental materials

Sterile *Salvia miltiorrhiza* plantlets were cultured on a Murashige and Skoog basal medium, as described previously (Yan and Wang, 2007). All chemicals were obtained from Sigma Chemical Co. (St. Louis, MO, USA). Solvents were of high-performance liquid chromatography (HPLC) grade. Standards of RA, Sal B, and JA were purchased from the National Institute for the Control of Pharmaceutical and Biological Products (Beijing, China). All were prepared as stock solutions in methanol and stored in the dark at ‒18°C. Primer pairs are listed in Supplementary Table 1 and Table 2.

### Construction of plant expression vectors and plant transformation

To construct the *SmbHLH37* overexpression vector, we amplified the full-length open reading frame (ORF) of *SmbHLH37* (GenBank Accession Number KP257470.1) with primers GV*SmbHLH37*-F/R, which introduced attB sites, and subsequently re-combined it into the pDONR207 vector (BP reaction Gateway®) according to the protocol from the Gateway manufacturer (Invitrogen, United States). The ENTRY vector pDONR207‒*SmbHLH37* was sequenced and inserted into the pEarleyGate 202 vector (Earley *et al.*, 2006) by an LR reaction (Gateway®) to generate the pEarleyGate 202‒*SmbHLH37* overexpression vector. *Agrobacterium*-mediated gene transfer was performed based on protocols established in our laboratory (Yan and Wang, 2007).

### Molecular detection of transgenic plantlets

To evaluate whether the overexpressing box had been integrated into the transgenic plant genome, we amplified the *CaMV35S* promoter from isolated genomic DNA, using previously published protocols (Yang *et al.*, 2017). Total RNA from the roots of *S. miltiorrhiza* transgenic lines was extracted and converted into cDNA. Gene expression was monitored via qRT-PCR, with housekeeping gene *SmUbiquitin* serving as an internal reference. Relative expression was analyzed according to the comparative 2^-ΔΔCt^ method (Livak and Schmittgen, 2001).

Based on the transcript levels of *SmbHLH37*, we conducted qRT-PCR analysis to determine the expression levels of key enzyme genes for the biosynthetic pathways of Sal B, JA, and anthocyanin. Each sample from a transgenic line was assayed with three independent biological replicates and three technical replicates.

### Determination of anthocyanin concentrations

Extraction and quantification of anthocyanins was performed in accordance with the protocols of Mano *et al* (2007), with minor modifications. 20 mg samples of powder from transgenic or wild type plants were extracted with 1 mL of acidic methanol (1% [v/v] HCl) for 1 h at 20°C, with moderate shaking (100 rpm). After centrifugation (12000 rpm, room temperature, 5 min), 0.7 mL of the supernatant was added to 0.7 mL of chloroform. Absorption of the extracts at wavelengths of 530 and 657 nm was determined photometrically (DU 640 Spectrophotometer, Beckman Instruments). Quantitation of anthocyanins was performed using the following equation: Q (anthocyanins)=(A530−0.25×A657)×M^−1^, where Q (anthocyanins) is the concentration of anthocyanins, A530 and A657 are the absorptions at the wavelengths indicated, and M is the dry weight (in grams) of the plant tissue used for extraction.

### Determination of phenolics and JA concentrations by LC/MS analysis

Roots were collected from two-month-old transgenic plantlets and air-dried at 20±2°C. The phenolic compounds were extracted and determined as described by Li *et al* (2018).

To determine the concentration of JA, we extracted JA using a modified protocol as described (Yang *et al.*, 2012). Approximately 0.1-g root samples were homogenized and added to 10 mL of cold extraction buffer (acetone: 50 mM citric acid, 7:3, v/v).

After this mixture was vortexed and then left to stand 30 min at 4°C, 10 mL of ethyl acetate was added before vortexing again. Following centrifugation at 5000 g for 10 min at 4°C, the supernatants were transferred to new 50-mL tubes and evaporated to dryness in a freeze dryer. The residue of each sample was re-suspended in 1 mL of 80% methanol (v/v) and sonicated for 10 min, then passed through a 0.22-μm organic filter. The extracts were loaded onto an Agela Cleanert SPE-NH2 (500 mg/6 mL); sonication and filtration steps were repeated. The combined supernatants were used for JA detection.

We determined the concentrations of JA in the plant samples by LC-QQQ-MS. Briefly, analyses were conducted using an Agilent 1260 HPLC system coupled to an Agilent 6460 QQQ LC-MS system equipped with a dual electrospray ion source operated in the negative mode. The extracts were separated on a Welch Ultimate XB-C18 column (2.1 × 150 mm, 3 μm). The chromatographic separation was performed over an 8-min analysis time, using a linear gradient of 85% to 50% A (0–6 min), 50% to 0% A (6–7 min), and 0% to 0% A (7–8 min). The flow rate of the gradient mobile phase was 0.4 mL/min, and the column temperature was 30°C. Conditions for mass spectrometry included a drying gas temperature of 300°C, drying gas flow of 10 L/min, nebulizer pressure of 45 psi, ion spray voltage of 3500 V, and sheath gas of 11 L/min, at a temperature of 350°C. Retention time was 6.9 min for JA. The precursor/product ion of JA was 209.1>59.1. The concentrations were quantified based on standard curves prepared with authentic reference standards.

### Bimolecular fluorescent complementation (BiFC)

The ORFs of *SmJAZ1*/*3*/*8* and *SmMYC2* without the termination codon were individually cloned into the pDONR207 vector through Gateway reactions and re-combined into the pEarleyGate202‒YC (YC) vector to generate YC‒*SmJAZ1/3/8* and YC-*SmMYC2*. Likewise, the ORF of *SmbHLH37* without the termination codon was inserted into the YC vector or pEarleyGate201‒YN (YN) to construct YC‒*SmbHLH37* and YN‒*SmbHLH37*. The YC and YN recombinant plasmids were mixed at equal densities before co-transformation.

The plasmids were transiently transformed into onion epidermis cells by particle bombardment (helium pressure, 1100 psi) with the PDS-1000/He system (Bio-Rad, CA, USA). After 24 h of incubation, those cells were stained with DAPI (Vector Labs, CA, USA) for 20 min and then observed using a Leica DM6000B microscope (Leica, Germany) with an excitation wavelength of 475 nm.

### Yeast two-hybrid (Y2H) assays

The full-length coding sequence of *SmbHLH37* was cloned into the pGADT7 or pGBKT7 vector, while those of *SmJAZ1/3/8* and *SmMYC2* were cloned into the pGADT7 vector. To test potential auto-activation of the prey, vectors of pGBKT7‒*SmbHLH37* and pGADT7 were co-transformed. Empty vectors of pGADT7 and pGBKT7 were also co-transformed as a negative control. The two types of recombinant vectors were co-transformed into yeast strain AH109 by the PEG/LiAC method (Zhou and Memelink, 2016). Interaction assays were performed according to manufacturer’s protocol for the Matchmaker Gold Yeast Two-Hybrid System (Clontech, USA), and Y2H images were taken on Day 5 of incubation.

### Yeast one-hybrid (Y1H) assays

The ORFs of *SmbHLH37* and *SmMYC2* were individually amplified by PCR using primers containing *Bam*HI and *Eco*RI restriction sites. They were fused to the GAL4 activation domain in vector pGADT7‒Rec2 (Clontech) to create the fusion proteins pGADT7‒*SmbHLH37* and pGADT7‒*SmMYC2*. The ~798-bp, ~1350-bp, and ~1146-bp promoter regions of *SmPAL1*, *SmTAT1*, and *SmCYP98A14*, respectively, were amplified and cloned into pHIS2 (Clontech). These recombinant vectors were co-transformed into yeast strain Y187 according to the reported protocol (Huang *et al.*, 2013). The transformed cells were cultured on an SD/-Leu/-Trp medium and then selected on an SD/-Leu/-Trp/-His medium supplemented with 60 mM 3-amino-1, 2, 4-triazole to examine any protein‒DNA interactions.

### Assay of transient transcriptional activity (TTA) in Nicotiana benthamiana

For assaying transient transcriptional activity, we amplified and cloned the ~798-bp, ~1350-bp, and ~1146-bp promoter regions of *SmPAL1*, *SmTAT1*, and *SmCYP98A14*, respectively, into the pGreenII 0800‒LUC (luciferase) vector (Hellens *et al.*, 2005) to generate our reporter construct. The full-length coding sequences of *SmMYC2* and *SmbHLH37* were inserted into the pGreenII62‒SK vector as the effector. Transient expression was monitored in *N. benthamiana* leaves according to the protocols of Sparkes *et al* (2006). After 3 d of infiltration, activities of firefly LUC and renillia luciferase (REN) were measured using a dual-luciferase reporter gene assay kit (Beyotime Biotechnology, China) and a GloMax 20/20 luminometer (Promega, USA). Relative LUC activity was calculated by normalizing it against REN activity.

## Results

### SmbHLH37 forms homodimer and interacts with SmJAZ3/8

*SmbHLH37* can be dramatically induced by exogenous MeJA (Zhang *et al.*, 2015). We previously showed that SmbHLH37 is most similar to AtJAM3 (Su *et al.*, 2017), which interacts with JAZs in *Arabidopsis* (Fonseca, 2014; Sasaki-Sekimoto *et al.*, 2014). To detect whether SmbHLH37 and SmJAZs could interact with each other in *S. miltiorrhiza*, we performed BiFC and Y2H assays. Our results demonstrated that SmbHLH37 interacts with SmJAZ3/8 (Fig. 1A, B). Among the JAZ proteins in *S. miltiorrhiza*, SmJAZ8 has been verified as being involved in the biosynthesis of salvianolic acids and tanshinones (Ge *et al.*, 2015; Pei *et al.*, 2018). The bHLH protein usually forms a homodimer or heterodimer to develop their function (Feller *et al.*, 2006). Our findings indicated that SmbHLH37 does form a homodimer (Fig. 1A, B). Moreover，SmbHLH37 do not interact with SmMYC2.

**Fig. 1.**
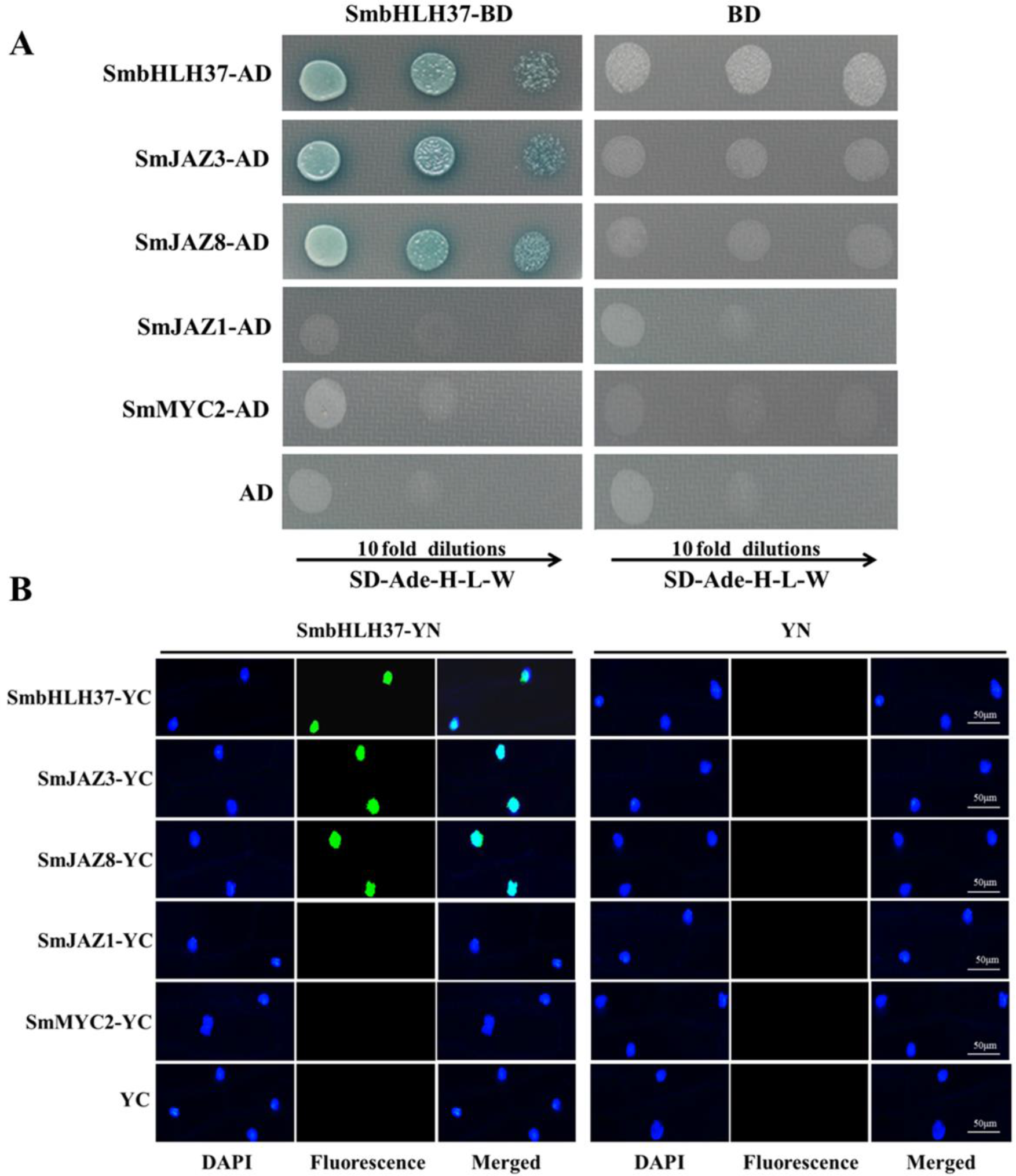
SmbHLH37 interacts with SmJAZ3, SmJAZ8, and SmbHLH37. (A) Yeast two-hybrid assay to detect interactions. SmJAZ1, SmJAZ3, SmJAZ8, SmMYC2, and SmbHLH37 were fused with activation domain (AD) while SmbHLH37 was simultaneously fused with DNA-binding domain (BD). Transformed yeast cells were grown on SD/-Ade/-Leu/-Trp/-His/X-α-gal media. Different rows represent individual dilutions of cells. (B) Bimolecular fluorescent complementation experiments in onion epidermis cells. SmJAZ1, SmJAZ3, SmJAZ8, SmMYC2, and SmbHLH37 were fused with C-terminal of fluorescin to produce SmJAZ1-YC, SmJAZ3-YC, SmJAZ8-YC, SmMYC2-YC, and SmbHLH37-YC, respectively. SmbHLH37 was fused with N-terminal of fluorescin to produce SmbHLH37-YN. Recombinant vectors were co-transformed with corresponding empty vectors as control. Nucleus was located after staining with DAPI.

### Overexpression of SmbHLH37 decreases endogenous JA concentrations and affects JA signal pathway

Using PCR amplifications, we confirmed that the transgenic plants indeed contained an expected 721-bp fragment of the CaMV 35S promoter (Supplementary Fig. S1A). Real-time quantitative PCR demonstrated that expression of *SmbHLH37* was highest in Lines OE-4 and OE-7 when compared with the non-transformed wild type (WT) (Supplementary Fig. S1B). Therefore, we chose those two lines for further analysis. JA is derived from a-linolenic acid and the biosynthesis pathway was shown in Fig. 2A. The transcript levels of genes encoding LOX (lipoxygenase), AOS (allene oxide synthase), AOC (allene oxide cyclase), and OPR3 (12-oxophytodienoic acid reductase) were significantly down-regulated in OE lines (Fig. 2B). We performed LC-MS to determine the concentrations of endogenous JA in fresh root samples from OE and WT lines. The MRM chromatograms, shown in Supplementary Fig. S2, revealed that those JA levels were significantly decreased in OE-4 and OE-7 when compared with the control (Fig. 2C). These results implied that *SmbHLH37* participates in JA biosynthesis in *S. miltiorrhiza*.

**Fig. 2.**
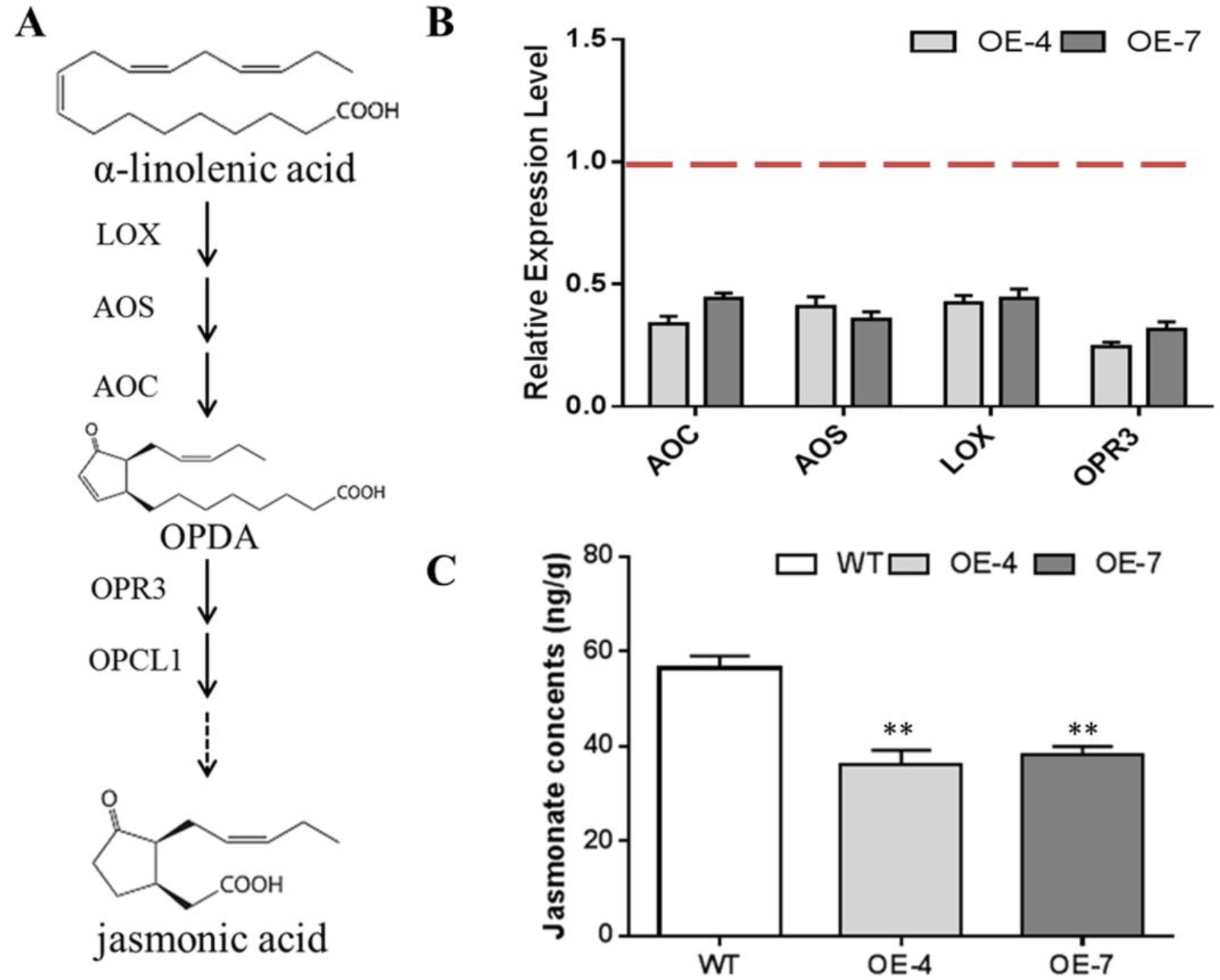
Effects of *SmbHLH37* overexpression on pathway of JA biosynthesis. (A) Pathway Enzymes: LOX, lipoxygenase; AOS, allene oxide synthase; AOC, allene oxide cyclase; OPR3, 12-oxophytodienoate reductase 3. (B) Relative expression levels of genes involved in JA biosynthesis pathway. Expression values in WT were set to ‘1’ (not shown). (C) Concentrations of JA in root extracts from *SmbHLH37*-overexpressing lines (OE) and wild type (WT). All data are means of 3 replicates, with error bars indicating SD; ∗∗; values are significantly different from WT at P <0.01.

We further examined the transcription changes of genes encoding JAZ proteins and MYC2, core factors in the JA signaling pathway. Our qRT-PCR results showed that overexpression of *SmbHLH37* significantly decreased the transcript levels of *SmJAZ1/3/8* and *SmMYC2* (Fig. 3).

**Fig. 3.**
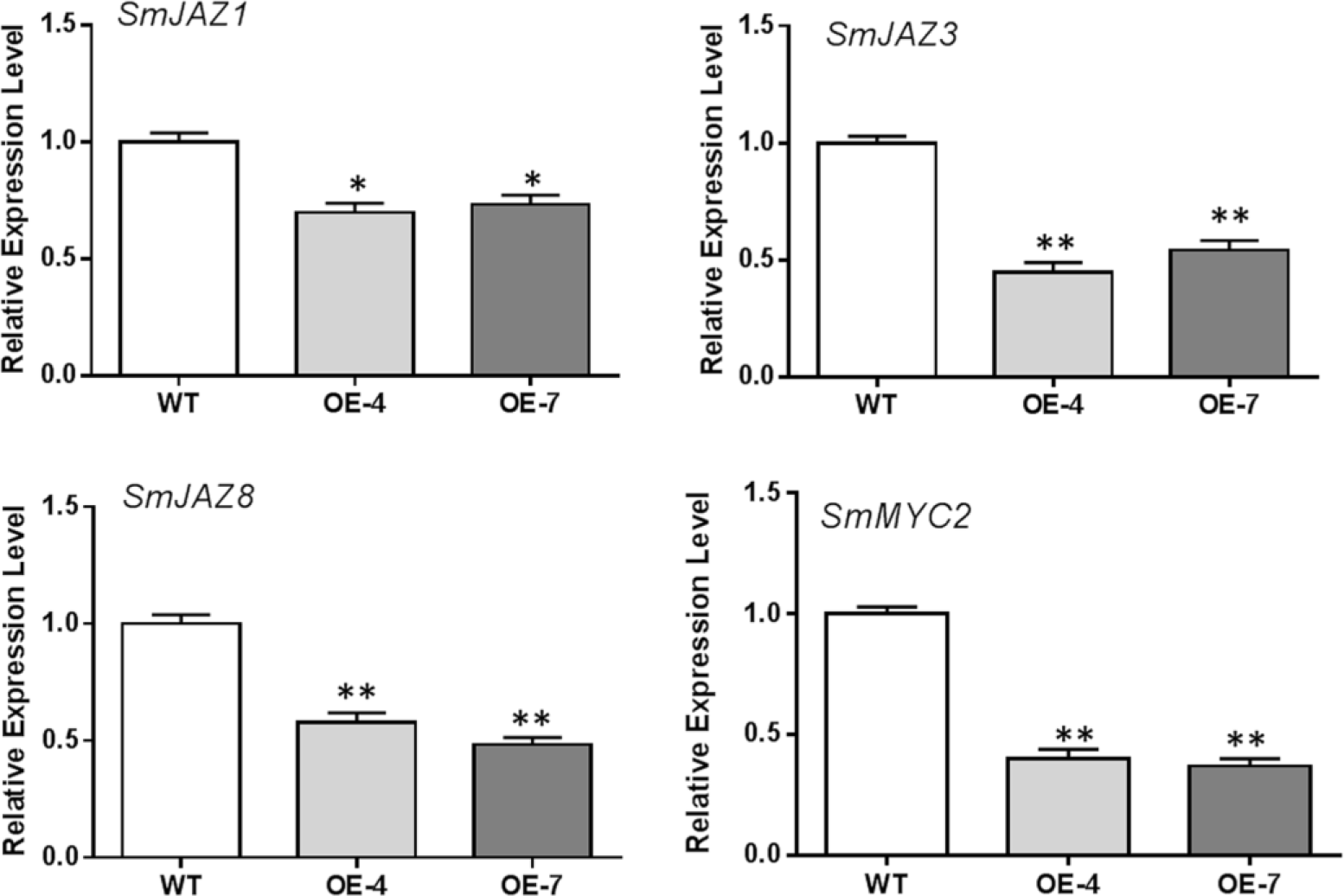
Results of qRT-PCR analysis on expression levels of *SmJAZ*s and *SmMYC2* in *SmbHLH37*-overexpressing lines (OE) and wild type (WT). All data are means of 3 replicates, with error bars indicating SD; ∗ and ∗∗, values are significantly different from WT at P <0.05 and P <0.01, respectively.

### SmbHLH37 negatively regulates anthocyanin biosynthetic pathway through transcriptional cascade

We tested whether this regulation of a transcriptional cascade by SmbHLH37 alters anthocyanin levels and found that concentrations of this pigment were significantly lower in OE-4 and OE-7 than in the WT (Fig. 4A, B). We also investigated the expression profiles of genes for anthocyanin biosynthesis, e.g., *CHS* (chalcone synthase), *F3′H* (flavonoid 3′-hydroxylase), *F3′5′H* (flavonoid 3′5′-hydroxylase), *FLS* (flavonol synthase), and *DFR* (dihydroflavonol 4-reductase). All were significantly down-regulated in OE lines, with *DFR* showing the largest fold-change (Fig. 4C).

**Fig. 4.**
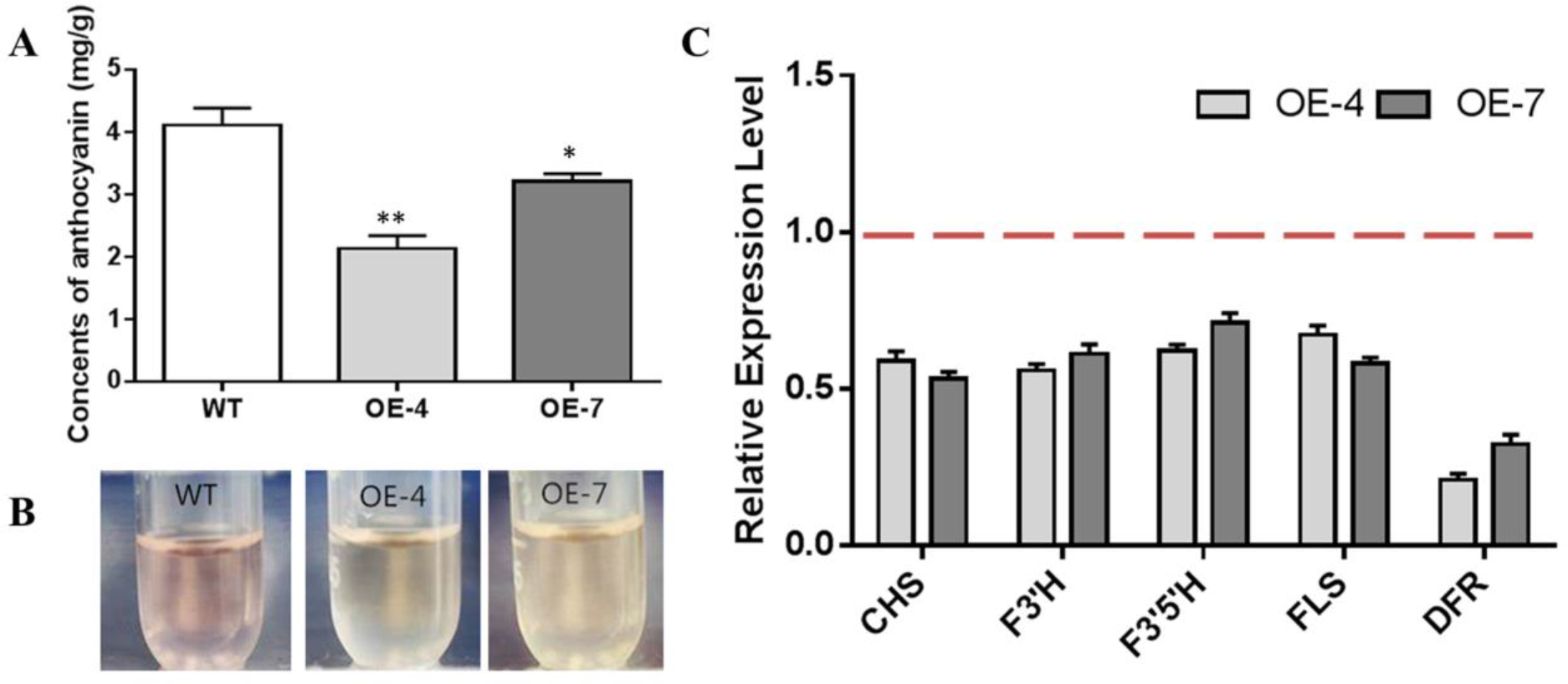
Effects of *SmbHLH37* overexpression on pathway for anthocyanin biosynthesis. (A) Concentrations of anthocyanin in roots of *SmbHLH37*-overexpressing lines (OE) and wild type (WT). (B) Color of root extracts. (C) Relative expression levels of genes involved in pathway. CHS, chalcone synthase; F3′H, flavonoid 3′-hydroxylase; F3′5′H, flavonoid 3′5′-hydroxylase; FLS, flavonol synthase; DFR, dihydroflavonol 4-reductase. Expression values in WT were set to ‘1’ (not shown). All data are means of 3 replicates, with error bars indicating SD; * and **, values are significantly different from WT at P <0.05 and P <0.01, respectively.

### Overexpression of SmbHLH37 decreases concentrations of phenolic acids

We predicted that the production of salvianolic acids would be decreased because of the decline in JA levels. To test this, we performed LC-MS to determine the concentrations of RA and Sal B. As shown by the MRM chromatograms in Supplementary Fig. S2, the results were consistent with our expectations, i.e., the levels of RA and Sal B were significantly declined in OE lines (respective reductions of 2.0-and 1.8-fold for RA and Sal B in OE-4; 1.7-and 1.6-fold for RA and Sal B in OE-7) when compared with the WT (Fig. 5). To evaluate how the expression of genes related to phenolic acid biosynthesis is influenced in transgenic lines, we monitored relative transcript levels for 11 enzyme genes in the WT and OE lines (Fig. 5). Expression of all tested genes was significantly decreased in OE plants. In particular, transcript levels of *RAS6* were decreased 6.0-and 5.1-fold in OE-4 and OE-7, respectively.

**Fig. 5.**
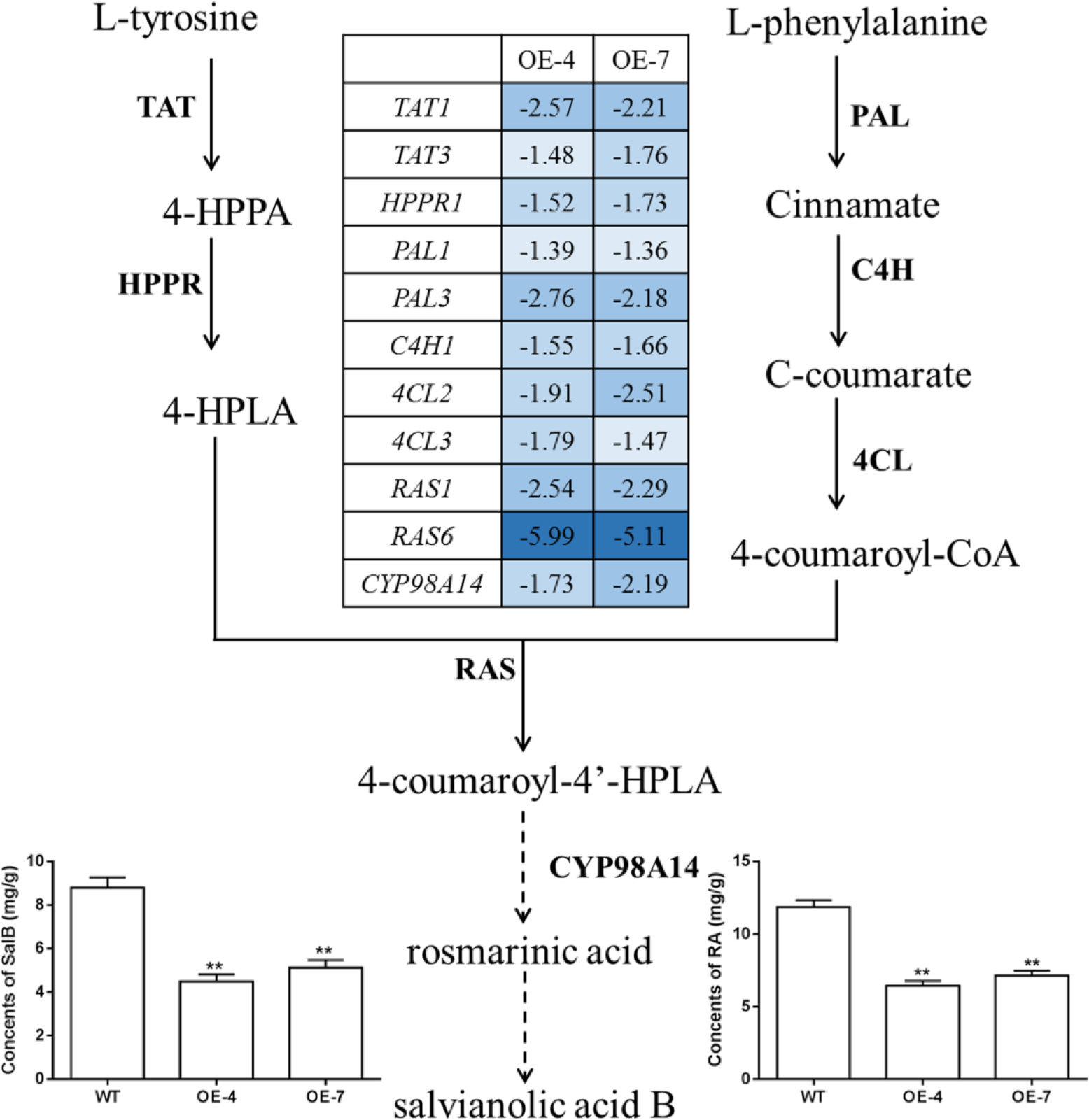
Effects of *SmbHLH37* overexpression on pathway for salvianolic acid biosynthesis. Enzymes: TAT, tyrosine aminotransferase; HPPR, hydroxyl phenylpyruvate reductase; PAL, phenylalanine ammonia lyase; C4H, cinnamate 4-hydroxylase; 4CL, hydroxycinnamate-CoA ligase; RAS, rosmarinic acid synthase; and CYP, cytochrome P450. Negative values in array indicate fold-change in *SmbHLH37*-overexpressing lines (O E-4 and OE-7) relative to wide type (WT). Bars show concentrations of salvianolic acid B (Sal B) and rosmarinic acid (RA) accumulated in roots of OEs and WT, determined by LC-MS. All data are means of 3 replicates, with error bars indicating SD; **, values are significantly different from WT at P <0.01.

### SmbHLH37 binds to and represses promoters of SmTAT1 and SmPAL1

The bHLH TFs function by binding to the E/G-box of the target gene promoter. Although 29 enzyme genes have been predicted to participate in phenolic acid biosynthesis in *S. miltiorrhiza* (Wang *et al.*, 2015), only a few have been verified as doing so, including *SmPAL1* (Song and Wang, 2011), *SmTAT1* (Xiao *et al.*, 2011), and *SmCYP98A14* (Di *et al.*, 2013). Each of them carries E/G-box sequences in its promoter (Fig. 6A). We speculated whether SmbHLH37 is directly involved in regulating the pathway of phenolic acid biosynthesis. Our results from the Y1H assay showed that SmbHLH37 directly binds to the promoters of *SmTAT1* and *SmPAL1* rather than *SmCYP98A14* in yeast (Fig. 6B).

**Fig. 6.**
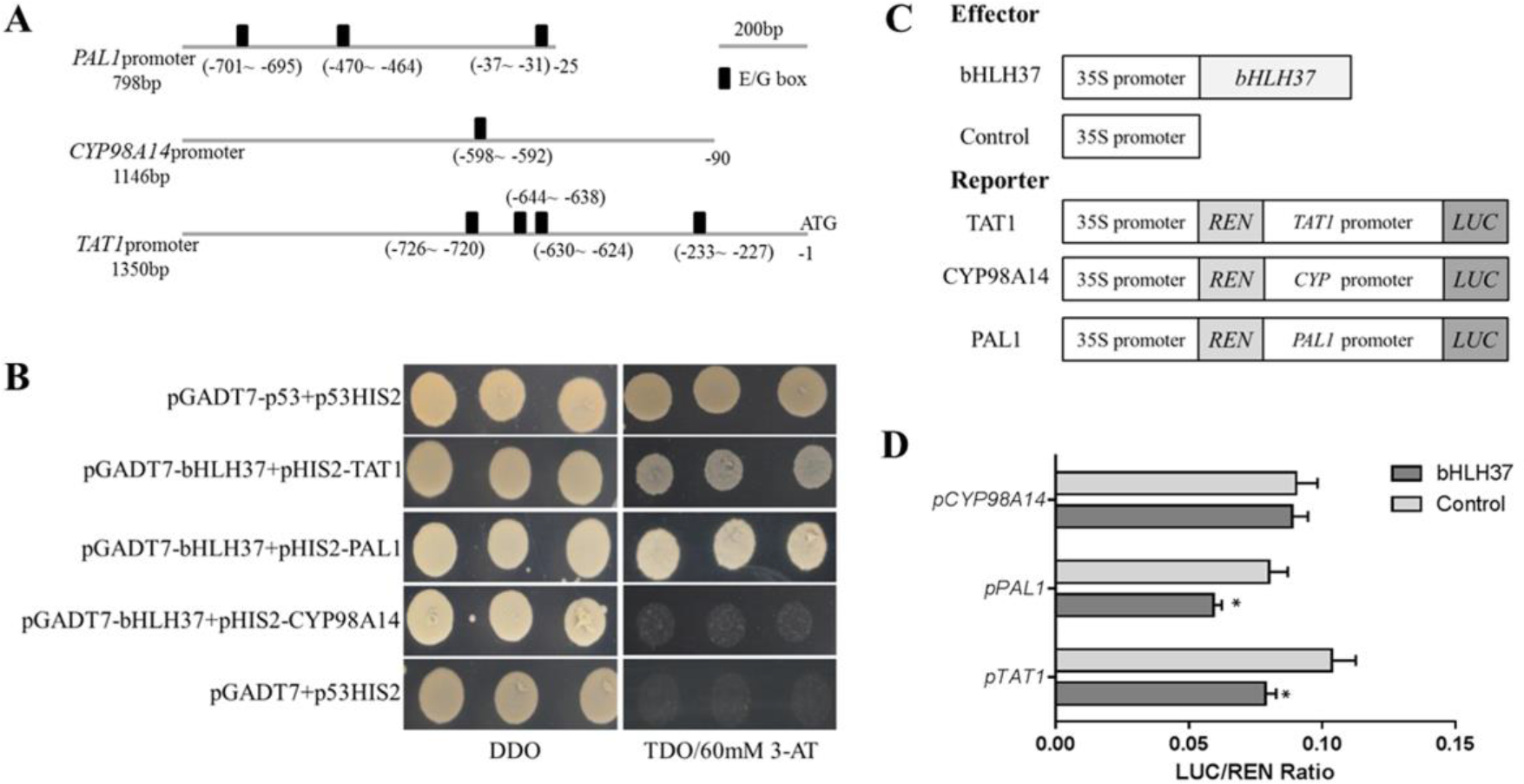
SmbHLH37 binds to and represses promoters of *SmTAT1* and *SmPAL1*. (A) E-box fragments of *SmTAT1*, *SmPAL1*, and *SmCYP98A14* promoters. (B) Yeast one-hybrid assay to detect interaction between SmbHLH37 and promoters of *SmTAT1*, *SmPAL1*, and *SmCYP98A14*. SmbHLH37 was fused to GAL4 activation domain (AD). Promoter regions of *SmTAT1*, *SmPAL1* and *SmCYP98A14* were cloned into pHIS2 to construct pHIS2-*SmTAT1*, pHIS2-*SmPAL1*, and pHIS2-*SmCYP98A14*, respectively. Recombinant vectors were co-transformed into yeast strain Y187, and transformed cells were cultured on SD/-Leu/-Trp medium (DDO), then selected on SD/-Leu/-Trp/-His medium (TDO) supplemented with 60 mM 3-amino-1, 2, 4-triazole (3-AT) to examine protein‒DNA interaction. The p53HIS2/pGADT7-p53 and p53HIS2/pGADT7 served as positive control and negative control, respectively. (C) Schematic diagram of constructs used in assays of transient transcriptional activity. (D) SmbHLH37 represses promoters of *SmTAT1* and *SmPAL1*. Effector SmbHLH37 was co-transformed with reporters *P*_TAT1_-*LUC*, *P*_*PAL1*_-*LUC*, and *P*_*CYP98A14*_-*LUC*. All data are means of 3 replicates, with error bars indicating SD; * and **, values are significantly different from WT at P <0.05 and P <0.01, respectively.

We then conducted an assay of transient transcriptional activity in *N. benthamiana* leaves. The promoter regions of *SmTAT1*, *SmPAL1*, and *SmCYP98A14* were fused individually with LUC to generate the reporter, and SmbHLH37, driven by the 35S promoter, was used as an effector (Fig. 6C). As showed in Fig. 6D, SmbHLH37 did bind to the promoter regions of *SmTAT1* and *SmPAL1* to repress the expression of LUC. The same was not true for *SmCYP98A14*. Therefore, these results demonstrated that SmbHLH37 directly binds to the promoter regions of *SmTAT1* and *SmPAL1* to repress their expression.

### SmMYC2 binds to and activates promoters of SmTAT1, SmPAL1, and SmCYP98A14

We have reported that overexpression of *SmMYC2* strongly increases the production of RA and Sal B, and those transcript levels of *SmTAT1* and *SmPAL1* are dramatically improved in *SmMYC2*-OE lines (Yang *et al.*, 2017). However, the molecular mechanism had not yet been characterized. Here, we performed Y1H assays and examined transient transcriptional activity to verify whether SmMYC2 directly binds to the promoter regions of these genes to activate their expression. Results from our Y1H assay showed that *Sm*MYC2 did bind to the promoter regions of *SmTAT1*, *SmPAL1*, and *SmCYP98A14* (Fig. 7A).

**Fig. 7.**
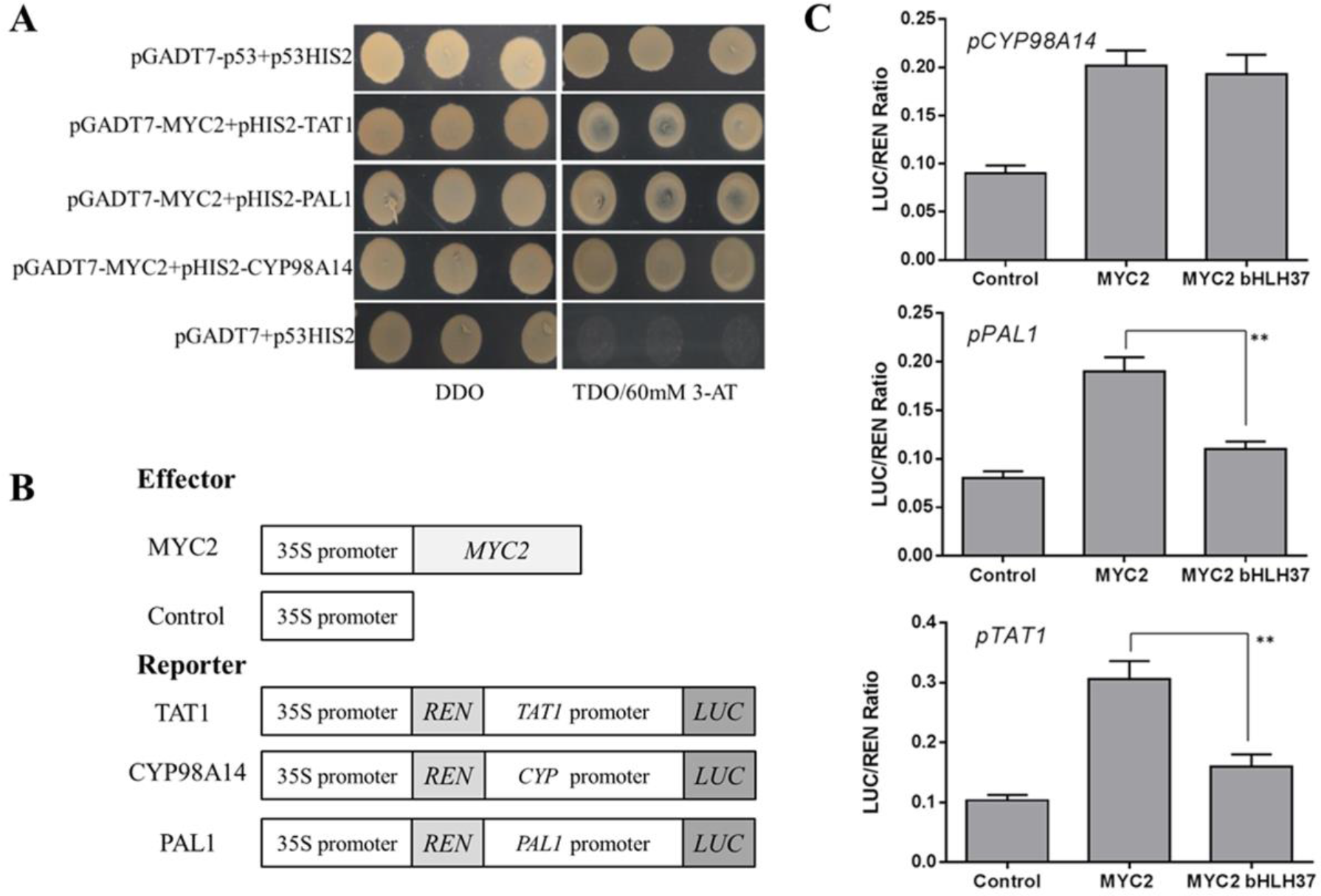
SmMYC2 binds to and activates promoters of *SmTAT1*, *SmPAL1*, and *SmCYP98A14*. (A) Yeast one-hybrid assay to detect interaction between SmMYC2 and promoters. SmMYC2 was fused to GAL4 activation domain (AD). Promoter regions of *SmTAT1*, *SmPAL1*, and *SmCYP98A14* were cloned into pHIS2 to construct pHIS2-*SmTAT1*, pHIS2-*SmPAL1*, and pHIS2-*SmCYP98A14*, respectively. Recombinant vectors were co-transformed into yeast strain Y187 and transformed cells were cultured on SD/-Leu/-Trp medium (DDO), then selected on SD/-Leu/-Trp/-His medium (TDO) supplemented with 60 mM 3-amino-1, 2, 4-triazole (3-AT) to examine protein‒DNA interaction. The p53HIS2/pGADT7-p53 and p53HIS2/pGADT7 served as positive control and negative control, respectively. (B) Schematic diagram of constructs used in assays of transient transcriptional activity. (C) Activation of *SmTAT1* and *SmPAL1* promoters by SmMYC2 is repressed by SmbHLH37. Effector SmMYC2, alone or together with SmbHLH37, was co-transformed with reporters *P*_*TAT1*_-*LUC*, *P*_*PAL1*_-*LUC*, and *P*_*CYP98A14*_-*LUC*. All data are means of 3 replicates, with error bars indicating SD; * and **, values are significantly different from WT at P <0.05 and P <0.01, respectively.

To conduct transient transcriptional activity analysis, SmMYC2 was used as an effector (Fig. 7B). SmMYC2 activated the expression of *SmTAT1*, *SmPAL1*, and *SmCYP98A14*, based on our data from the assay of transient transcriptional activity (Fig. 7C). We also learned that SmbHLH37 can repress SmMYC2-activated LUC expression, as driven by the promoters of *SmTAT1* and *SmPAL1* (Fig. 7C). Together, these findings suggested that SmbHLH37 antagonizes transcription activator SmMYC2 in the Sal B biosynthesis pathway.

## Discussion

Jasmonates are widely distributed in the plant kingdom (Browse, 2009). They are derived from a-linolenic acid and the biosynthetic enzymes consist of LOX, AOS, AOC, and OPR (Wasternack, 2007). The JAM1/2/3, members of the bHLHs IIId subfamily in *A. thaliana*, have redundant functions that negatively regulate the JA metabolic pathway (Nakata *et al.*, 2013; Nakata and Ohme-Takagi, 2013). We previously reported that SmbHLH37 is most closely associated with AtJAM3 and belongs to the IIId subfamily (Su *et al.*, 2017). Here, overexpression of *SmbHLH37* significantly decreased the level of endogenous JA by repressing the transcripts of *LOX*, *AOC*, *AOS*, and *OPR3*. This indicated that SmbHLH37 is involved in JA biosynthesis in *S. miltiorrhiza*.

Application of exogenous MeJA is an effective way to improve the yields of secondary metabolites. Earlier research showed that JA signaling has a role in the biosynthesis of salvianolic acids and tanshinones (Xiao *et al.*, 2009; Zhang *et al.*, 2011b; Pei *et al.*, 2018). Expression of genes in the salvianolic acid and tanshinone biosynthetic pathways is increased significantly after MeJA treatment (Ge *et al.*, 2015; Pei *et al.*, 2018). Our results also indicated that overexpression of *SmbHLH37* significantly decreased RA and Sal B concentrations. Such accumulation profiles were consistent with the expression profiles of all the tested genes involved in Sal B biosynthesis. We previously proposed that *SmbHLH37* helps modulate tanshinone biosynthesis because it is up-regulated by MeJA treatment and is more highly expressed in the roots than in any other organs (Zhang *et al.*, 2015). We also detected tanshinone IIA and cryptotanshinone but found no significant differences in amounts between control plants and *SmbHLH37*-OE lines (data not shown).

Activation of JA signaling can also improve the accumulation of anthocyanin in *S. miltiorrhiza* (Ge *et al.*, 2015). Here, overexpression of *SmbHLH37* significantly decreased the levels of anthocyanin as well as the expression of genes in its biosynthetic pathway. One gene, *DFR*, has a vital role in anthocyanin production (Lim *et al.*, 2016), and we noted that it had the greatest fold-change among the five genes tested here. Therefore, overexpression of *SmbHLH37* repressed overall the biosynthetic pathways for JA, anthocyanin, and salvianolic acids, which is contrary to the activation of JA signaling.

MYC2 is a core TF in the plant response to jasmonates, inducing JA-mediated responses such as wounding, inhibition of root growth, JA and anthocyanin biosynthesis, and adaptations to oxidative stress (Dombrecht *et al.*, 2007). The JAZ proteins directly interact with MYC2 and inhibit its activity, meaning that they function as repressors of the JA pathway (Chini *et al.*, 2007; Thines *et al.*, 2007; Seo *et al.*, 2011; Song *et al.*, 2011). In *S. miltiorrhiza*, the SmJAZs have proven to be negative regulators of salvianolic acid and tanshinone biosynthesis (Ge *et al.*, 2015; Shi *et al.*, 2016; Pei *et al.*, 2018). In contrast, the orthologs of MYC2 act as positive regulators (Zhou *et al.*, 2016; Yang *et al.*, 2017). Although overexpression of *SmMYC2* increases the production of phenolic acids in *S. miltiorrhiza* (Yang *et al.*, 2017), the responsible molecular mechanism is still unclear.

The bHLH TFs function by binding to the E/G box of the target gene promoters (Shoji and Hashimoto, 2011). Transcriptomic and qRT-PCR analyses of *SmMYC2*-OE and control plants of *S. miltiorrhiza* have shown that transcript levels for *SmPAL1* and *SmTAT1* are increased by 367.1-fold and 110-fold, respectively, in the transgenics (Yang *et al.*, 2017). Both genes contain the E/G-box sequences in their promoters. Our Y1H and transient transcriptional activity assays with tobacco leaves also demonstrated that SmMYC2 directly binds to the promoters of *SmPAL1* and *SmTAT1* to activate their expression. Previous electronic mobility shift assays have shown that SmMYC2a and SmMYC2b bind with the E-box within the *SmCYP98A14* promoter *in vitro* (Zhou *et al.*, 2016). We also confirmed here that SmMYC2 up-regulates the expression of *SmCYP98A14* by binding to its promoter in yeast. Our analysis indicated that the sequence of *SmMYC2a* is almost completely consistent with that of *SmMYC2*. Therefore, we speculate that they are the same gene.

In *Arabidopsis*, JAM1/2/3 function as transcription repressors to antagonize the transcription activator MYC2 by binding to its target sequences (Song et al., 2013; Qi *et al.*, 2015b). Our results from Y1H and transient transcriptional activity assays showed that SmbHLH37 represses *SmPAL1* and *SmTAT1* by binding to their promoters. Moreover, we found that SmbHLH37 employs antagonistic regulation with SmMYC2 by binding to the promoters of the same target genes. These results are consistent with the relationship described between JAM1/2/3 and MYC2 in *Arabidopsis* (Song *et al.*, 2013; Qi *et al.*, 2015b).

Based on our results and previous reports, we propose a model to illustrate the JA-induced accumulation of salvianolic acids (Fig. 8). In it, we confirm that SmbHLH37 regulates such accumulations in *S. miltiorrhiza* by engineering the biosynthetic pathway genes. That protein also shows antagonistic regulation with *Sm*MYC2 because they bind to the promoters of the same target genes. Jasmonate induces the degradation of JAZ proteins, thereby releasing SmMYC2 and SmbHLH37. The former binds to and activates the promoters of genes involved in salvianolic acid biosynthesis (e.g., *SmTAT1*, *SmPAL1*, and *SmCYP98A14*), ultimately promoting the accumulation of those salvianolic acids. Meanwhile, SmbHLH37 represses these genes and antagonizes this accumulation that is activated by SmMYC2. Both *SmbHLH37* and *SmJAZ*s are more highly expressed in *SmMYC2*-OE lines than in the control (Su *et al.*, 2017; Yang *et al.*, 2017). In contrast, we found here that expression of *SmMYC2* and *SmJAZ*s was lower in *SmbHLH37*-OE lines than in the WT. These data suggest that SmMYC2 activates SmJAZs and SmbHLH37, while SmbHLH37 suppresses SmMYC2 and SmJAZs. Although more research is needed on the relationships among SmJAZs, SmMYC2, and SmbHLH37, we speculate that over-expressing SmMYC2 and silencing SmbHLH37 simultaneously is a promising genetic engineering strategy to dramatically enhance concentrations of salvianolic acids.

**Fig. 8.**
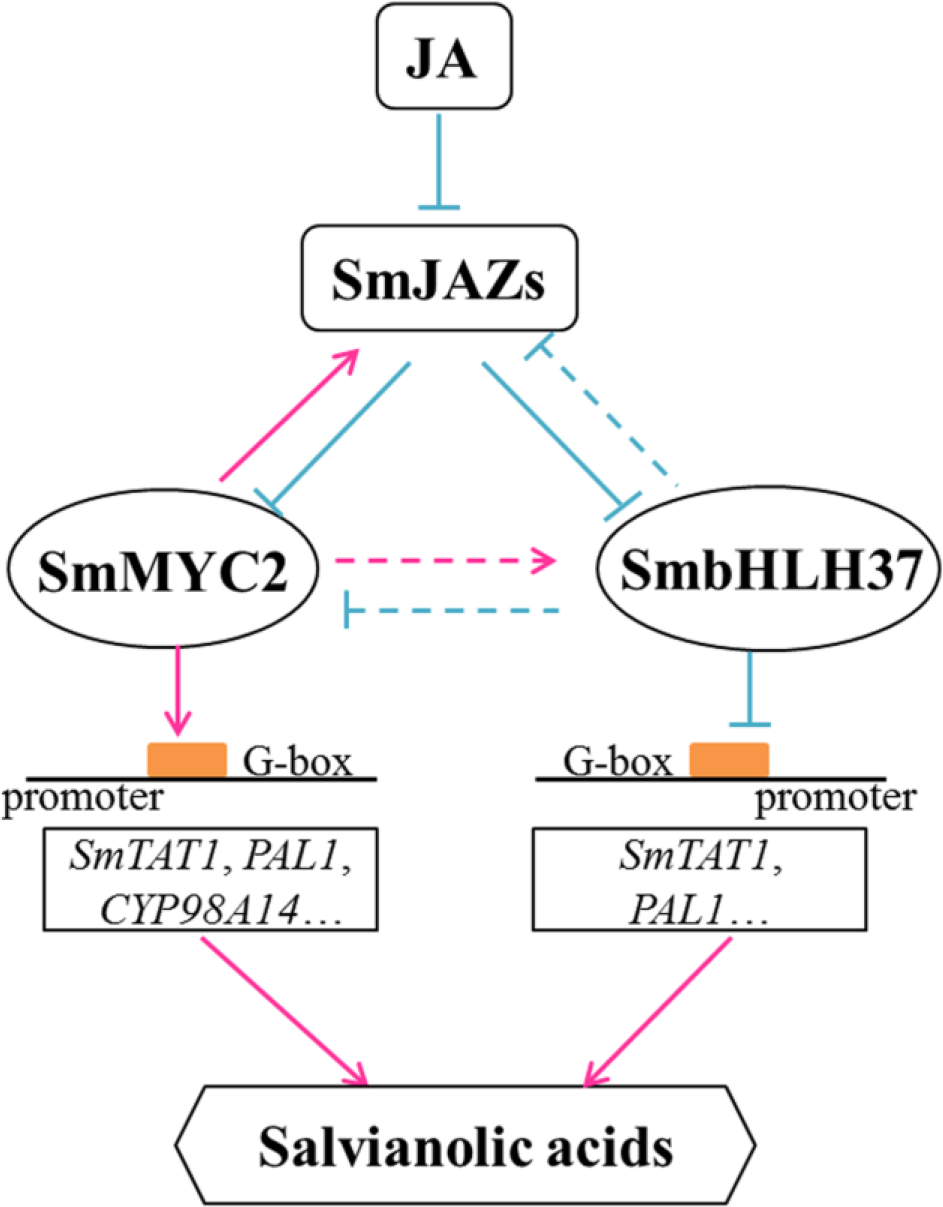
Model illustrating regulation of salvianolic acid biosynthesis by SmbHLH37. Upon perception of JA, JAZ proteins are targeted for degradation. SmbHLH37 and SmMYC2 are then released to regulate, antagonistically or coordinately, their target genes (e.g., *SmTAT1* and *SmPAL1*), which further modulates accumulation of salvianolic acids. SmbHLH37 acts as transcription repressor of JA signaling in *Salvia miltiorrhiza*.

## Supplementary data

**Fig. S1.** Detection of *SmbHLH37*-overexpressing transgenic lines of *Salvia miltiorrhiza*.

**Fig. S1.** MRM maps of JA standard and samples.

**Figure 3** MRM maps of RA and Sal B standards and samples.

**Table**S**1.** Primers used for vector construction.

**Table**S**2.** Primers used for quantitative real-time PCR.

## Acknowledgements

This work was funded by the National Natural Science Foundation of China (31870276 and 31670299); the Natural Science Foundation of Shaanxi Province, China (2018JZ3001); the National Key Technologies R & D Program for Modernization of Traditional Chinese Medicine (2017YFC1701300); Major Project of Shaanxi Province, China (2017ZDXM-SF-005); and Fundamental Research Funds for the Central Universities (GK201706004).

## Abbreviations

*SmbHLH37*: *Salvia miltiorrhiza* basic helix-loop-helix 37
*SmTAT1*: *Salvia miltiorrhiza* tyrosine aminotransferase 1
*SmPAL1*: *Salvia miltiorrhiza* phenylalanine ammonialyase 1
*SmCYP98A14*: *Salvia miltiorrhiza* cytochrome P450 monooxygenase 98A14
Sal B: salvianolic acid B

## References

Browse J. 2009. Jasmonate passes muster: a receptor and targets for the defense hormone. Annual Review of Plant Biology 60,183–205.

Carretero-Paulet L, Galstyan A, Roig-Villanova I, Martínez-García JF, Bilbao-Castro JR, Robertson DL. 2010. Genome-wide classification and evolutionary analysis of the bHLH family of transcription factors in Arabidopsis, poplar, rice, moss, and algae. Plant Physiology 153, 1398–1412.

Chico JM, Chini A, Fonseca S, Solano R. 2008. JAZ repressors set the rhythm in jasmonate signaling. Current Opinion in Plant Biology 11,486–494.

Chini A, Fonseca S, Fernãn G, Adie B, Chico JM, Lorenzo O, Garcã-A-Casado G, Lã³Pez-Vidriero I, Lozano FM, Ponce MR. 2007. The JAZ family of repressors is the missing link in jasmonate signalling. Nature 448,666–671.

Di P, Zhang L, Chen J, Tan H, Ying X, Xin D, Xun Z, Chen W. 2013. ^13^C Tracer Reveals Phenolic Acids Biosynthesis in Hairy Root Cultures of Salvia miltiorrhiza. Acs Chemical Biology 8, 1537–1548.

Dombrecht B, Xue G, Sprague S, Kirkegaard J, Ross J, Reid J, Fitt G, Sewelam N, Schenk P, Manners J.2007. MYC2 differentially modulates diverse jasmonate-dependent functions in Arabidopsis. Plant Cell 19,2225–2245.

Earley KW, Haag JR, Pontes O, Opper K, Juehne T, Song K, Pikaard CS. 2006. Gateway-compatible vectors for plant functional genomics and proteomics. Plant Journal 45,616–629.

Ezer D, Shepherd SJ, Brestovitsky A, Dickinson P, Cortijo S, Charoensawan V, Box MS, Biswas S, Jaeger K, Wigge PA.2017. The G-box transcriptional regulatory code in Arabidopsis. Plant Physiology 175,628–640.

Farmer EE.2007. Jasmonate perception machines. Nature. 448, 659–660.

Feller A, Hernandez JM, Grotewold E.2006. An ACT-like domain participates in the dimerization of several plant basic-helix-loop-helix transcription factors. Journal of Biological Chemistry 281,28964–28974.

Fonseca S, Fernándezcalvo P, Fernández GM, Díezdíaz M, Gimenezibanez S, Lópezvidriero I, Godoy M, Fernándezbarbero G, Leene JV, Jaeger GD. 2014. bHLH003, bHLH013 and bHLH017 are new targets of JAZ repressors negatively regulating JA responses. Plos One 9, e86182.

Ge Q, Zhang Y, Hua WP, Wu YC, Jin XX, Song SH, Wang ZZ.2015. Combination of transcriptomic and metabolomic analyses reveals a JAZ repressor in the jasmonate signaling pathway of Salvia miltiorrhiza. Scientific Reports 5, 14048.

Goossens J, Mertens J, Goossens A. 2017. Role and functioning of bHLH transcription factors in jasmonate signalling. Journal of Experimental Botany 68, 1333–1347.

Guo Y, Li Y, Xue L, Severino RP, Gao S, Niu J, Qin LP, Zhang D, Brömme D. 2014. Salvia miltiorrhiza: An ancient Chinese herbal medicine as a source for anti-osteoporotic drugs. Journal of Ethnopharmacology 155, 1401–1416.

Han JY, Fan JY, Horie Y, Miura S, Cui DH, Ishii H, Hibi T, Tsuneki H, Kimura I. 2008. Ameliorating effects of compounds derived from Salvia miltiorrhiza root extract on microcirculatory disturbance and target organ injury by ischemia and reperfusion. Pharmacology & Therapeutics 117, 280–295.

Hellens RP, Allan AC, Friel EN, Bolitho K, Grafton K, Templeton MD, Karunairetnam S, Gleave AP, Laing WA. 2005. Transient expression vectors for functional genomics, quantification of promoter activity and RNA silencing in plants. Plant Methods 1, 13–13.

Hichri I, Barrieu F, Bogs J, Kappel C, Delrot S, Lauvergeat V. 2011. Recent advances in the transcriptional regulation of the flavonoid biosynthetic pathway. Journal of Experimental Botany 62, 2465–2483.

Huang XS, Wang W, Zhang Q, Liu JH.2013. A basic helix-loop-helix transcription factor, PtrbHLH, of Poncirus trifoliata confers cold tolerance and modulates peroxidase-mediated scavenging of hydrogen peroxide. Plant Physiology 162,1178–1194.

Katsir L, HS C, AJ K, GA H. 2008. Jasmonate signaling: a conserved mechanism of hormone sensing. Current Opinion in Plant Biology 11, 428–435.

Lim SH, You MK, Kim DH, Kim JK, Lee JY, Ha SH.2016. RNAi-mediated suppression of dihydroflavonol 4-reductase in tobacco allows fine-tuning of flower color and flux through the flavonoid biosynthetic pathway. Plant Physiology and Biochemistry 109,482–490.

Livak KJ, Schmittgen TD.2001. Analysis of relative gene expression data using real-time quantitative PCR and the 2^-ΔΔCt^ method. Method 25,402–408.

Li SS, Wu YC, Kuang J, Wang HQ, Du TZ, Huang YY, Zhang Y, Cao XY, Wang ZZ.2018. SmMYB111 is a key factor to phenolic acid biosynthesis and interacts with both SmTTG1 and SmbHLH51 in Salvia miltiorrhiza. Journal of Agricultural & Food Chemistry 66,8069–8078.

Lorenzo O, Chico JM, Sánchezserrano JJ, Solano R. 2004. JASMONATE-INSENSITIVE1encodes a MYC transcription factor essential to discriminate between different jasmonate-regulated defense responses in Arabidopsis. Plant Cell 16, 1938–1950.

Ma P, Liu J, Osbourn A, Dong J, Liang Z.2015. Regulation and metabolic engineering of tanshinone biosynthesis. RSC Advances. 5, 18137–18144.

Ma P, Liu J, Zhang C, Liang Z. 2013. Regulation of water-soluble phenolic acid biosynthesis in Salvia miltiorrhiza Bunge. Applied Biochemistry & Biotechnology 170, 1253–1262.

Mano H, Ogasawara F, Sato K, Higo H, Minobe Y.2007. Isolation of a regulatory gene of anthocyanin biosynthesis in tuberous roots of purple-fleshed sweet potato. Plant Physiology 143, 1252–1268.

Nakata M, Mitsuda N, Herde M, Koo AJ, Moreno JE, Suzuki K, Howe GA, Ohme-Takagi M.2013. A bHLH-type transcription factor, ABA-INDUCIBLE BHLH-TYPE TRANSCRIPTION FACTOR/JA-ASSOCIATED MYC2-LIKE1, acts as a repressor to negatively regulate jasmonate signaling in Arabidopsis. Plant Cell 25,1641–1656.

Nakata M, Ohme-Takagi M.2013. Two bHLH-type transcription factors, JA-ASSOCIATED MYC2-LIKE2 and JAM3, are transcriptional repressors and affect male fertility. Plant Signaling & Behavior 8,e26473.

Pei T, Ma P, Ding K, Liu S, Jia Y, Ru M, Dong J, Liang Z. 2018. SmJAZ8 acts as a core repressor regulating JA-induced biosynthesis of salvianolic acids and tanshinones in Salvia miltiorrhiza hairy roots. Journal of Experimental Botany 69, 1663–1678.

Pires N, Dolan L.2010. Origin and diversification of basic-helix-loop-helix proteins in plants. Molecular Biology & Evolution 27, 862–874.

Qi T, Huang H, Song S, Xie D. 2015a. Regulation of Jasmonate-Mediated Stamen Development and Seed Production by a bHLH-MYB Complex in Arabidopsis. Plant Cell 27,1620–1633.

Qi T, Wang J, Huang H, Liu B, Gao H, Liu Y, Song S, Xie D.2015b. Regulation of jasmonate-induced leaf senescence by antagonism between bHLH subgroup IIIe and IIId factors in Arabidopsis. Plant Cell 27,1634–1649.

Sasaki-sekimoto Y, Saito H, Masuda S, Shirasu K, Ohta H.2014. Comprehensive analysis of protein interactions between JAZ proteins and bHLH transcription factors that negatively regulate jasmonate signaling. Plant Signaling & Behavior 9, e27639.

Seo JS, Joo J, Kim MJ, Kim YK, Nahm BH, Song SI, Cheong JJ, Lee JS, Kim JK, Choi YD.2011. OsbHLH148, a basic helix-loop-helix protein, interacts with OsJAZ proteins in a jasmonate signaling pathway leading to drought tolerance in rice. Plant Journal 65, 907–921.

Sheard LB, Xu T, Mao H, John W, Gili BN, Hinds TR, Yuichi K, Fong-Fu H, Michal S, John B.2010. Jasmonate perception by inositol phosphate-potentiated COI1-JAZ co-receptor. Nature 468, 400–405.

Shi M, Zhou W, Zhang J, Huang S, Wang H, Kai G.2016. Methyl jasmonate induction of tanshinone biosynthesis in Salvia miltiorrhiza hairy roots is mediated by JASMONATE ZIM-DOMAIN repressor proteins. Scientific Reports 6,20919.

Shoji T, Hashimoto T.2011. Tobacco MYC2 regulates jasmonate-inducible nicotine biosynthesis genes directly and by way of the NIC2-locus ERF genes. Plant Cell Physiology 52, 1117–1130.

Song J, Wang ZZ. 2011. RNAi-mediated suppression of the phenylalanine ammonia-lyase gene in Salvia miltiorrhiza causes abnormal phenotypes and a reduction in rosmarinic acid biosynthesis. Journal of Plant Research 124, 183–192.

Song S, Qi T, Fan M, Zhang X, Gao H, Huang H, Wu D, Guo H, Xie D. 2013. The bHLH subgroup IIId factors negatively regulate jasmonate-mediated plant defense and development. Plos Genetics 9, e1003653.

Song S, Qi T, Huang H, Ren Q, Wu D, Chang C, Peng W, Liu Y, Peng J, Xie D. 2011. The Jasmonate-ZIM domain proteins interact with the R2R3-MYB transcription factors MYB21 and MYB24 to affect Jasmonate-regulated stamen development in Arabidopsis. Plant Cell 23, 1000–1013.

Sparkes IA, Runions J, Kearns A, Hawes C. 2006. Rapid, transient expression of fluorescent fusion proteins in tobacco plants and generation of stably transformed plants. Nature Protocols 1, 2019–2025.

Su CY, Ming QL, Khalid R, Han T, Qin LP. 2015. Salvia miltiorrhiza: Traditional medicinal uses, chemistry, and pharmacology. Chinese Journal of Natural Medicines 13, 163–182.

Su J, Yang N, Wang X, Du T, Li S, LU Zhang T, Cao X. 2017. Cloning, subcellular localization and expression analysis of SmbHLH37 gene in Salvia miltiorrhiza. Journal of Agricultural Biotechnology 25, 884–892.

Thines B, Katsir L, Melotto M, Niu Y, Mandaokar A, Liu G, Nomura K, He SY, Howe GA, Browse J.2007. JAZ repressor proteins are targets of the SCF(COI1) complex during jasmonate signalling. Nature 448, 661–665.

Todd AT, Liu E, Polvi SL, Pammett RT, Page JE.2010. A functional genomics screen identifies diverse transcription factors that regulate alkaloid biosynthesis in Nicotiana benthamiana. Plant Journal 62,589–600.

Tsai M, Lin YL, Huang YT.2010. Effects of salvianolic acids on oxidative stress and hepatic fibrosis in rats. Toxicology and Applied Pharmacology 242, 155–164.

Wang B, Sun W, Li Q, Li Y, Luo H, Song J, Sun C, Qian J, Zhu Y, Hayward A.2015. Genome-wide identification of phenolic acid biosynthetic genes in Salvia miltiorrhiza. Planta 241, 711–725.

Wang D, Song Y, Chen Y, Yao W, Li Z, Liu W, Yue S, Wang Z.2013. Metabolic pools of phenolic acids in Salvia miltiorrhiza are enhanced by co-expression of Antirrhinum majus Delila and Rosea1 transcription factors. Biochemical Engineering Journal 74, 115–120.

Wasternack C.2007. Jasmonates: an update on biosynthesis, signal transduction and action in plant stress response, growth and development. Annals of Botany 100, 681–697.

Xiao Y, Gao SH, Peng D, Chen JF, Chen WS, Zhang L.2009. Methyl jasmonate dramatically enhances the accumulation of phenolic acids in Salvia miltiorrhiza hairy root cultures. Physiologia Plantarum 137,1–9.

Xiao Y, Zhang L, Gao S, Saechao S, Di P, Chen J, Chen W.2011. The c4h, tat, hppr and hppd genes prompted engineering of rosmarinic acid biosynthetic pathway in Salvia miltiorrhiza hairy root cultures. Plos One 6, e29713.

Yan YP, Wang ZZ,2007. Genetic transformation of the medicinal plant Salvia miltiorrhiza by Agrobacterium tumefaciens-mediated method. Plant Cell Tissue & Organ Culture 88, 175–184.

Yang DH, Hettenhausen C, Baldwin IT, Wu J.2012. Silencing Nicotiana attenuata calcium-dependent protein kinases, CDPK4 and CDPK5, strongly up-regulates wound-and herbivory-induced jasmonic acid accumulations. Plant Physiology 159, 1591–1607.

Yang N, Zhou W, Su J, Wang X, Li L, Wang L, Cao X, Wang Z.2017. Overexpression of SmMYC2 increases the production of phenolic acids in Salvia miltiorrhiza. Frontiers in Plant Science 8,1804.

Zeng M, Pan L, Qi S, Cao Y, Zhu H, Guo L, Zhou J.2013. Systematic review of recent advances in pharmacokinetics of four classical Chinese medicines used for the treatment of cerebrovascular disease. Fitoterapia 88, 50–75.

Zhang H, Hedhili S, Montiel G, Zhang Y, Chatel G, Pré M, Gantet P, Memelink J.2011. The basic helix-loop-helix transcription factor CrMYC2 controls the jasmonate-responsive expression of the ORCA genes that regulate alkaloid biosynthesis in Catharanthus roseus. Plant Journal 67, 61–71.

Zhang S, Liu Y, Shen S, Liang Z, Yang D. 2011. Effects of elicitors on accumulation of phenolic acids and tanshinones in Salvia miltiorrhiza hairy root. China Journal of Chinese Materia Medica 36, 1269–1274.

Zhang X, Luo H, Xu Z, Zhu Y, Ji A, Song J, Chen S.2015. Genome-wide characterisation and analysis of bHLH transcription factors related to tanshinone biosynthesis in Salvia miltiorrhiza. Scientific Reports 5, 11244.

Zhang Y, Yan YP, Wang ZZ. 2010. The Arabidopsis PAP1 transcription factor plays an important role in the enrichment of phenolic acids in Salvia miltiorrhiza. Journal of Agricultural & Food Chemistry 58, 12168–12175.

Zhang Y, Yan YP, Wu YC, Hua WP, Chen C, Ge Q, Wang ZZ.2014. Pathway engineering for phenolic acid accumulations in Salvia miltiorrhiza by combinational genetic manipulation. Metabolic Engineering 21,71–80.

Zhao GR, Zhang HM, Ye TX, Xiang ZJ, Yuan YJ, Guo ZX, Zhao LB. 2008. Characterization of the radical scavenging and antioxidant activities of danshensu and salvianolic acid B. Food & Chemical Toxicology 46, 73–81.

Zhou M, Memelink J.2016. Jasmonate-responsive transcription factors regulating plant secondary metabolism. Biotechnology Advances 34, 441–449.

Zhou Y, Sun W, Chen J, Tan H, Xiao Y, Li Q, Ji Q, Gao S, Chen L, Chen S. 2016. SmMYC2a and SmMYC2b played similar but irreplaceable roles in regulating the biosynthesis of tanshinones and phenolic acids in Salvia miltiorrhiza. Scientific Reports 6, 22852.

